# Phage-Mediated Iron Acquisition by *Pseudomonas aeruginosa*

**DOI:** 10.64898/2026.01.26.700923

**Authors:** Arya Khosravi, Aditi Gupta, Maryam Hajfathalian, Elizabeth B. Burgener, Saumel P. Rodriguez, Carlos E. Milla, Paul L. Bollyky

## Abstract

Iron acquisition is critical to bacterial growth and pathogenesis. Here, we describe a previously unrecognized mechanism by which the major human pathogen *Pseudomonas aeruginosa* acquires iron through a unique partnership with bacteriophages (phages). Pf is a filamentous phage that infects *P. aeruginosa* and is associated with chronic infections. We reveal that Pf contributes to *P. aeruginosa* pathogenesis by promoting iron uptake. We demonstrate that Pf phage is highly induced under iron-deplete growth conditions and that once induced, Pf virions directly bind and locally concentrate free iron. Pf-bound iron is more efficiently utilized by *P. aeruginosa* than unbound iron, enhancing bacterial growth. We further demonstrate that Pf-mediated iron acquisition depends on type IV pili, which facilitate Pf attachment and confers strain selective uptake of phage-bound iron, providing a competitive fitness advantage in polymicrobial settings. Together, these findings identify an unrecognized role for filamentous phages in mediating iron acquisition and reveal a novel phage–bacterium partnership that operates through selective kin cooperation.

## Introduction

*Pseudomonas aeruginosa* is a leading cause of treatment-refractory chronic lung and wound infections, resulting in frequent and prolonged hospitalizations, substantially higher healthcare costs and increased mortality^1,2^. These infections are especially devastating in individuals with underlying lung disease, including people with cystic fibrosis (pwCF), where *P. aeruginosa* infections result in progressive lung injury and premature death^3,4^. Once chronic infections are established, eradication is severely limited by the pathogen’s intrinsic and acquired resistance mechanisms. A key factor contributing to disease persistence is the formation of biofilms, multicellular bacterial communities encased in an extracellular matrix which shield bacteria from antimicrobial therapies^56,78,9^. These clinical challenges highlight an urgent need to better define the pathophysiology that enables *P. aeruginosa* to establish and maintain persistent infection.

In investigating unique factors driving chronic bacterial infections, we have identified Pf bacteriophage (phage), a filamentous Inovirus that infects *P. aeruginosa*, as a novel virulence factor. Its non-lytic replication cycle enables a mutualistic partnership in which Pf engages in multiple pathways to enhance *P. aeruginosa* virulence and persistence^10^. These effects translate into Pf phage promoting more severe disease in *P. aeruginosa* wound and lung infection models^11–13^. Clinically, the presence of Pf phage in pwCF with chronic *P. aeruginosa* lung infections is associated with greater infection chronicity and declines in lung function^14,15^. Similarly, among individuals with chronic *P. aeruginosa* wound infections, Pf positive strains are associated with delayed healing and increased wound progression^11,12^. Given these findings, further investigating the impact of Pf on *P. aeruginosa* virulence, including understanding the factors that regulate Pf expression, may provide key insights into disease pathophysiology and highlight new avenues for therapeutic intervention.

A defining feature of Pf phage is its ability to form highly-ordered crystalline networks that strengthen biofilm architecture^16,17^. These tactoid structures are formed by lateral bundling of Pf filaments with nematic ordering, which arise through depletion attraction forces^17,18^. However, filamentous phage can also form crosslinked networks in the presence of multivalent metal cations that demonstrate distinct physical properties compared to tactoid aggregates^17,19,20^. We previously demonstrated Pf phage forms aggregate polymer in the presence of ferric iron and that this polymer formation results in the loss of free iron^27^. Further, we showed that Pf phage treatment of *Aspergillus fumigates,* an opportunistic fungal pathogen, results in growth inhibition which was reversed following supplementation with free iron^27^. While this work provides some insight into the biological relevance of phage-metal assemblies and suggest a role of Pf phage on iron homeostasis, critical questions remain including how does Pf phage affect iron acquisition by *P. aeruginosa* itself, and what is its impact on bacterial fitness in the polymicrobial conditions characteristic of chronic infections.

Iron is a critical determinant of microbial pathogenesis. Iron availability influences the growth, biofilm development, and virulence of *P. aeruginosa* ^21,22^. As such, *P. aeruginosa* deploys multiple iron acquisition strategies. This includes uptake of ferric iron by siderophores pyoverdine and pyochelin, direct uptake of ferrous iron by the FeoB transport system, and heme utilization by HasR and PhuR receptors^23–25^. As excess iron is toxic, these uptake systems are tightly regulated in response to iron availability^24,25^. Under iron-depleted conditions, the extracytoplasmic function (ECF) sigma factor PvdS activates transcription of pyoverdine biosynthesis genes and other iron-starvation response genes^24,32^. Conversely, under iron-replete conditions, the transcriptional repressor Fur complexes with ferrous iron, forming homodimers that bind to promoter regions and represses iron acquisition genes, including PvdS^24^. While these systems have been well characterized in planktonic culture, iron acquisition during chronic, biofilm-associated infections remains poorly understood. Importantly, chronic infection is associated with a loss of canonical siderophore systems such as pyoverdine, suggesting selective for iron acquisition strategies that are tailored to the biofilm and host environment^26^.

Here, we have asked whether Pf phage impacts iron acquisition by its host bacterium. As Pf phage forms structural units within *P. aeruginosa* biofilms and binds cationic metal ions, including iron, we hypothesized that Pf may serve to sequester iron within the biofilm microenvironment during chronic infection.

## Results

### Pf phage binds free iron

To test the iron-binding capacity of Pf phage, we first validated its potential to bind and concentrate free iron. We demonstrate that Pf4 phage isolated from *P. aeruginosa* strain PA01 forms a gel polymer when mixed with iron (**Fig 1a,b**). Closer evaluation with transmission electron microscopy of purified Pf4 revealed individual phage filaments that when mixed with iron from a crosslinking aggregate network (**Fig 1c,d**). As iron serves as an ionic bridge to promote Pf crosslinking, these gel structures may concentrate free iron within the biofilm matrix. Indeed, as we previously shown, in the presence of Pf phage, there is reduced recovery of free, unbound iron in solution (**Fig 1d**)^27^. Together, these data demonstrate the capacity of Pf phage to directly bind free iron leading to the question of whether this sequestered iron remains bioavailable to *P. aeruginosa*.

**Figure 1.**
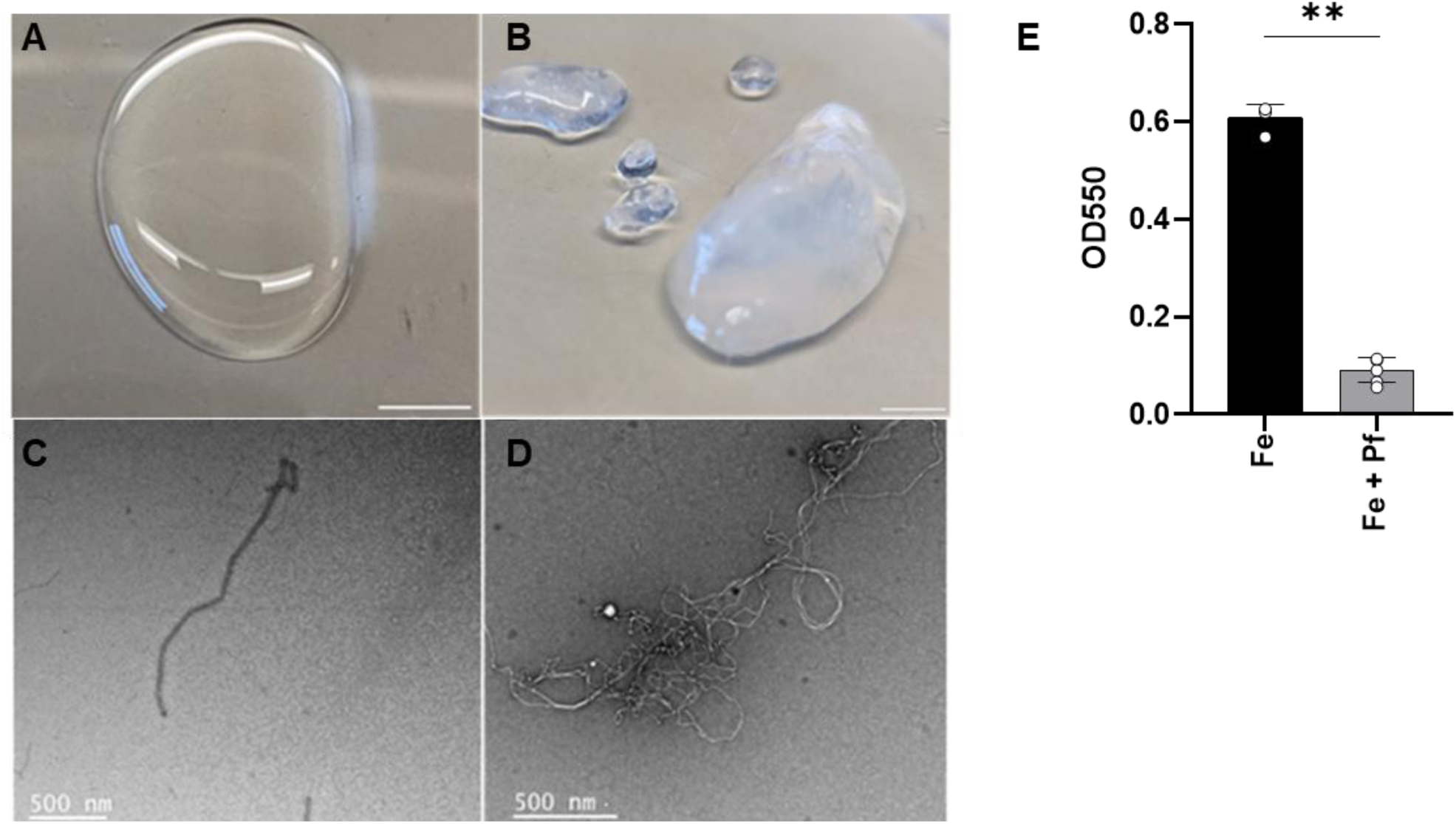
Pf phage binds free iron. Purified Pf4 phage mixed with deionize water vehicle control (A) or 100 µM ferric chloride (B) Bars represent 1 cm. Transmission electron microscopy imaging of Pf phage treated with vehicle control (C) or ferric chloride (D). Ferric chloride mixed with saline or Pf. E) Free iron availability measured by ferrozine colorimetric assay. ** p<0.01 by Student’s T-test.

### Pf phage promotes iron uptake and growth

To assess if Pf-bound iron is acquired by *P. aeruginosa*, purified Pf4 phage mixed with or without iron, was supplemented to cultures of PA14 and growth as an indirect assessment of iron uptake. PA14 was used rather than PAO1, as its growth is not impaired when supplemented with free Pf4 phage. Concentrations of supplemented ferric iron ranged between 1 and 100 µM, reflecting physiologic levels of iron measured in the sputum of healthy individuals and pwCF^28^. Growth was measured by optical density over time as well as direct quantification of live bacteria.

Iron supplementation enhanced PA14 growth, relative to media control while Pf4 phage alone had no effect (**Fig. 2a–c and S2a,b**). Notably, supplementation with Pf4 and iron together resulted in greater growth than equivalent concentrations of iron alone (**Fig. 2a–c and S2a,b**). This effect was specific to iron, as other Pf-polymerizing metal cations did not enhance Pseudomonas growth (**Fig. S2c,d**). We next assessed whether Pf enhanced ferrous-iron dependent growth. Ferric iron is the dominant form of iron under aerobic conditions; however, in individuals with chronic lung disease, microenvironments with reduced oxygen tension develop, creating conditions where ferrous iron becomes more available. Siderophores are required to acquire poorly soluble ferric iron, whereas ferrous iron is soluble and is imported directly via the FeoB transport system. Interestingly, we demonstrate that, in contrast to siderophores, Pf enhances both ferric and ferrous-dependent Pseudomonas growth (**Fig S2e,f**). Moreover, this Pf4-dependent enhancement of iron-mediated growth was conserved across multiple clinical *P. aeruginosa* isolates collected from pwCF, which exhibited increased growth when supplemented with Pf4 and iron compared to iron alone (**Fig. S2g-j**), demonstrating the beneficial impact of Pf4 on iron extends to multiple *P. aeruginosa* strains.

**Figure 2.**
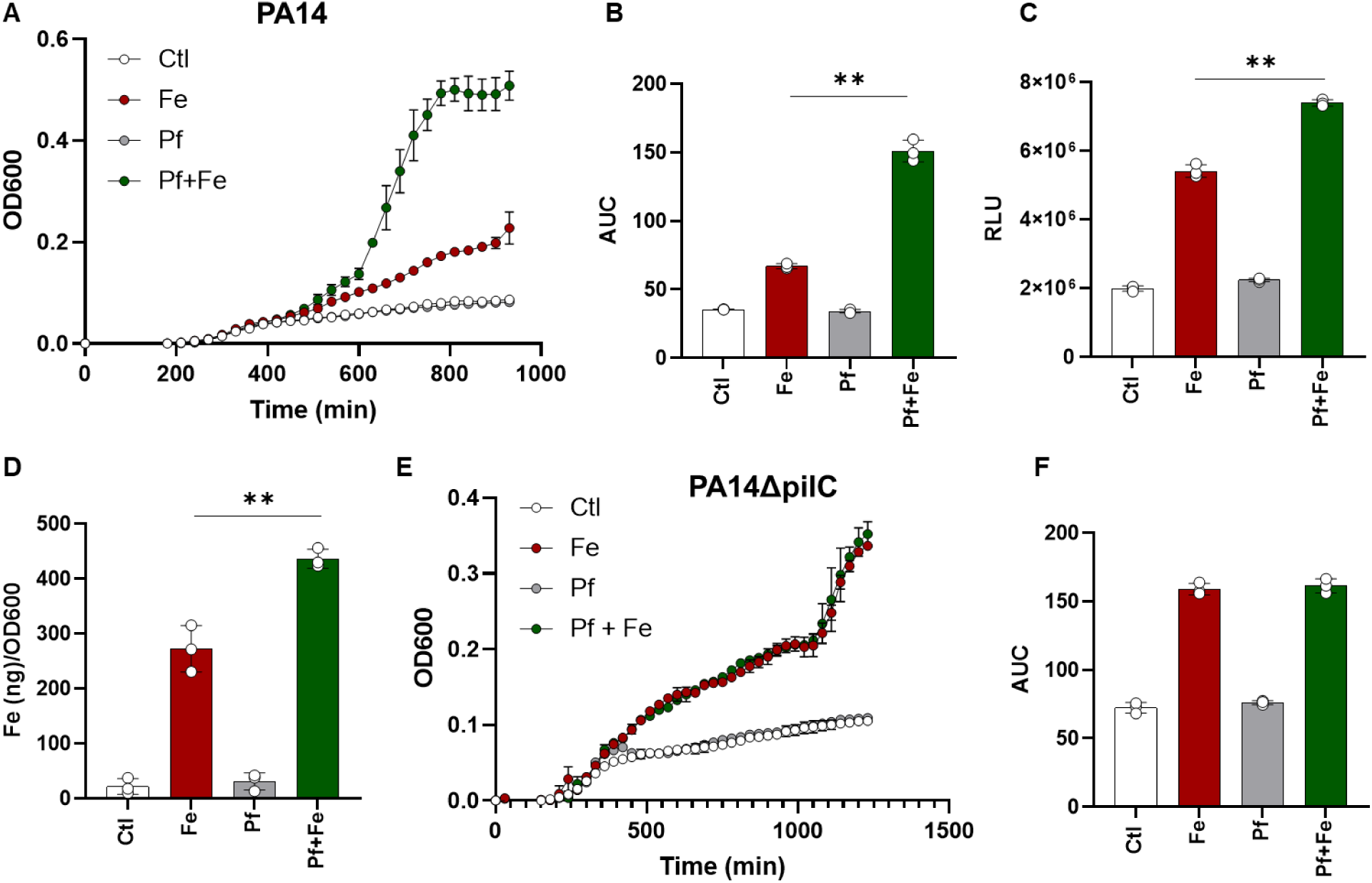
Pf promotes iron-dependent growth. PA14 is grown in M9 minimal media (Control, Ctl) or with 100 µM ferric chloride with or without Pf4 (1x1011 PFU/ml final concentration). Growth is measured over time (A) and area under the curve (AUC) is quantitated (B). C) Viable bacteria is quantitated by BacTiter-Glo and reported as relative light units (RLU). PA14 is grown in M9 minimal media with or without iron or Pf supplementation as above for eight hours then total bacterial iron is quantitated by Inductively Coupled Plasma-Mass Spectrometry (ICP-MS) and normalized to bacterial growth (D). PA14ΔpilC is grown in M9 media with or without iron or Pf4 phage supplementation and growth is measured over time (E-F). * p < 0.05 ** p < 0.01 by one-way ANOVA.

Because bacterial growth is an indirect measure of iron uptake, we directly quantified intracellular iron using inductively coupled plasma mass spectrometry (ICP-MS). PA14 was grown in minimal media with or without iron and/or Pf4 phage supplementation. After 8 hours, bacterial pellets were harvested, washed to remove extracellular iron, and total intracellular iron was quantified and normalized to bacterial growth. As expected, minimal intracellular iron was detected in cells grown in minimal media or with Pf4 phage alone (**Fig. 2d**). In contrast, co-supplementation with Pf4 phage and iron significantly increased intracellular iron levels compared with iron alone (**Fig. 2d**). These findings demonstrate that Pf phage promotes iron acquisition, consistent with the enhanced growth observed.

To define the mechanism underlying Pf-mediated iron uptake, we examined the role of type IV pili, which Pf phage binds via its minor coat protein to facilitate uptake into *P. aeruginosa*^10^. We observed that isogenic mutants lacking type IV pili failed to exhibit Pf4-dependent enhancement of iron-driven growth (**Fig. 2e–f**). Together, these data demonstrate that Pf phage binds iron and promotes its delivery into *P. aeruginosa* in a type IV pili–dependent manner, resulting in enhanced bacterial growth.

### Iron depletion triggers Pf expression

Iron uptake factors, including siderophores pyoverdine and pyochelin, are selectively expressed during iron-deplete growth conditions^21^. As Pf phage enhances iron uptake, we next asked whether iron availability might reciprocally regulate Pf expression.

To determine if iron impacts Pf induction, endogenous expression of Pf4 phage by PAO1 was measured in iron-deplete M9 minimal medial with or without iron supplementation. As expected, growth in media with less than 100 nM of iron resulted in impaired bacterial growth as well as enhanced expression of pyoverdine, an iron scavenging siderophore that is induced under iron-deplete growth conditions (**Fig 3a,b**). Under these low iron concentration conditions, Pf production is at its peak (**Fig 3c,d**). Conversely, increasing iron availability resulted in a dose-dependent inhibition of Pf phage, with a 50-fold reduction of Pf detected at 10 and 100 µM of iron. Indirect assessment of Pf expression, through quantification of Pf replicative form (RF), the excised prophage genome that is in active replication, also revealed enhanced Pf expression under iron limited growth conditions (**Fig S3a**). Of note, concentrations of iron that resulted in inhibition of Pf phage also resulted in loss of pyoverdine expression, supporting the notion that regulation of Pf expression is linked to iron uptake systems (**Fig 3b-d**). Further, inhibition of Pf was specific to iron, as other essential metals, including manganese, zinc, copper and nickel had no effect on Pf expression (**Fig S3b**). These studies demonstrate that ferric iron, the predominant iron species during aerobic growth, suppresses Pf expression under planktonic conditions.

**Figure 3.**
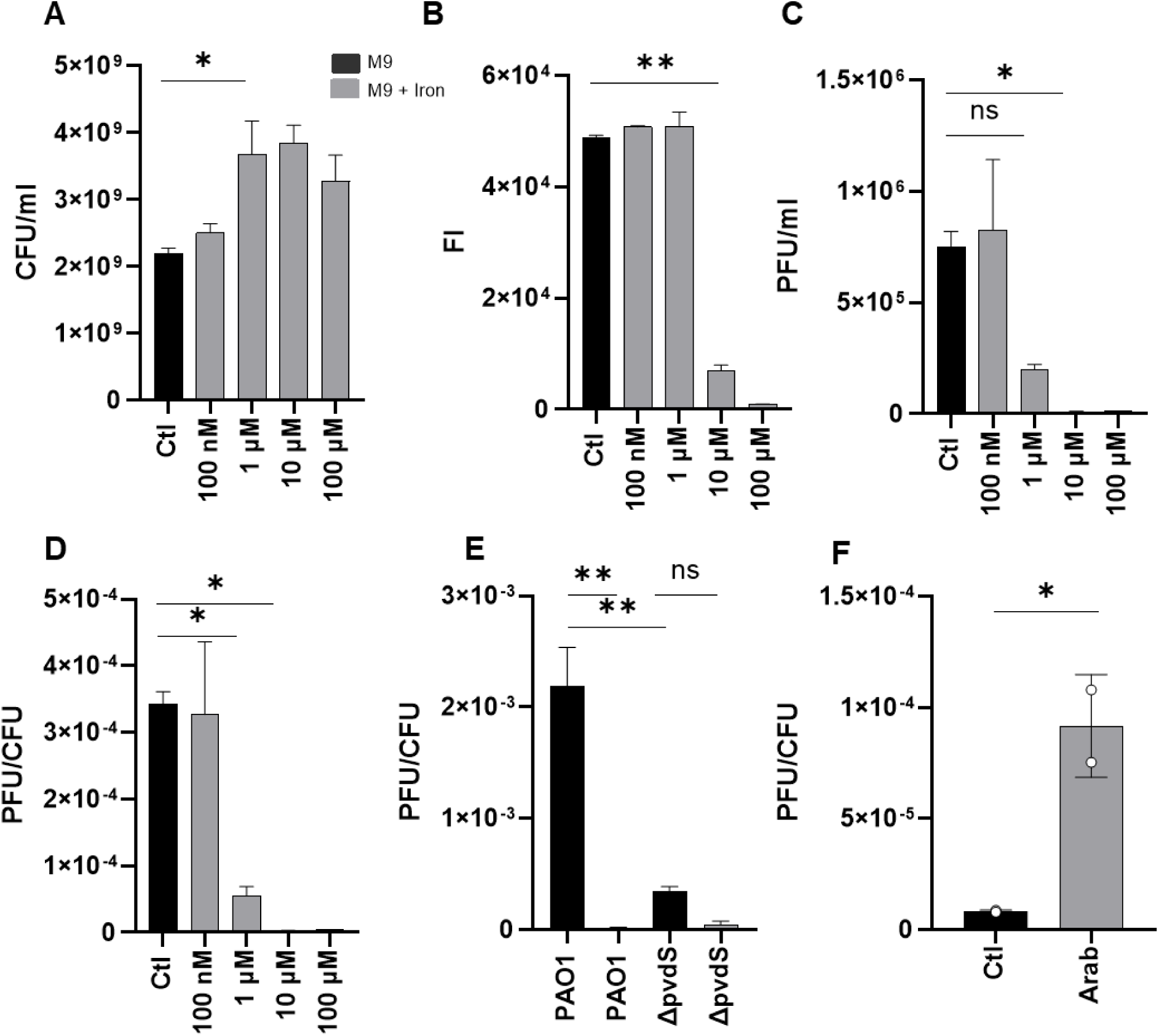
Iron-depletion drives Pf phage expression. A) PAO1 is grown in M9 minimal media without iron supplementation (Ctl) or increasing concentrations of iron. Bacterial growth (A), pyoverdine expression (B), Pf plaque formation (C), and Pf expression relative to bacteria (D) is measured at 24 hours of growth. E) PAO1 or PAO1ΔpvdS grown in M9 minimal media with or without 100 µM FeCl_3_ supplementation and Pf expression relative to bacterial growth is measured at 24 hours. F) PAO1 containing pvdS cloned under a arabinose (Arab)-inducible promoter treated with vehicle control or 0.5% arabinose. Pf expression relative to bacterial growth is measured following induction with Arab or vehicle control (Ctl). * p < 0.05 ** p < 0.01 by one-way ANOVA or Student’s T-test.

We next asked whether iron similarly regulates Pf expression in environments that more closely model chronic infection. Consistent with this, we found that ferrous iron and heme, which become major iron sources during chronic infection, also inhibit Pf expression (**Fig. S3c,d**). Chronic infections are further characterized by the formation of bacterial biofilms, a growth state in which Pf expression is markedly elevated^13,29^. This increase is driven in part by disruption of the phage repressor genes *pf4r* and *mvaU*, resulting in a 100–10,000-fold increase in phage production^30,31^. In addition to these known regulatory mechanisms, we now show that iron depletion further enhances Pf production during biofilm growth (**Fig. S3e**).

Finally, to determine whether iron-mediated inhibition of Pf is unique to PAO1, we examined clinical *P. aeruginosa* strains isolated from pwCF with chronic Pseudomonas infections and found that iron similarly suppresses Pf expression across multiple strains (**Fig. S3f**).

### PvdS drives Pf expression during iron-deplete conditions

A key regulator of the iron starvation response in *P. aeruginosa* is PvdS, an ECF sigma factor which drives gene expression necessary for iron uptake. GeneChip expression analysis of PAO1 identified 118 genes with greater than 5-fold induction during iron starvation, of which, close to a quarter of genes required PvdS for expression^32^.

To investigate the impact of PvdS on Pf expression, a pvdS isogenic deletion mutant PAO1ΔpvdS was grown in iron-deplete and -replete media. While wild-type PAO1 demonstrates elevated Pf expression during iron-deplete growth conditions, this induction of Pf phage was absent in PAO1ΔpvdS (**Fig 3e**). We observed that under iron-replete growth conditions, during which PvdS expression is inhibited, both wild-type and pvdS mutant demonstrate reduced Pf production (**Fig 3e**).

We then evaluated whether pyoverdine, which forms a regulatory loop with PvdS, is necessary to drive Pf expression. PvdS is bound and inhibited by pyoverdine receptor FpvA^33^. Upon binding of iron-bound pyoverdine (ferripyoverdine), FpvA undergoes proteolysis resulting in PvdS release which mediates gene expression including induction of pyoverdine^23,34^. We observed that similar to pvdS deletion strain, PAO1ΔpvdA, which lacks expression of pyoverdine, has impaired Pf induction during iron-deplete conditions (**Fig S3g**)^33^.

To further evaluate the role of PvdS in regulating Pf expression, pvdS was expressed under an arabinose inducible promoter. This construct was validated by demonstrating arabinose-induction of pvdS lead to enhanced expression of pyoverdine compared to uninduced control (**Fig S3h**). Induction of pvdS was sufficient to drive Pf production (**Fig 3f**).

Together, these studies reveal that PvdS signaling pathway drives Pf expression under iron-deplete growth conditions.

### Pf phage selectively promotes iron uptake by *P. aeruginosa* in polymicrobial growth conditions

Chronic infections are typically polymicrobial, comprising bacteria from multiple genera and taxonomic orders. We therefore sought to determine whether Pf-bound iron functions as a shared community resource or whether kin discrimination restricts its benefits to *Pseudomonas* isolates.

In contrast to enhanced growth seen with PA14 and clinical Pseudomonas isolates, supplementation of Pf phage with iron had minimal to no effect relative to iron alone among other Gram-negative pathogens including *Klebsiella pneumoniae, Klebsiella aeruginosa* and *Serratia marcescens* subspecies *marcescens*. (**Fig 4a and S4a-d**). To test whether Pf phage selectively deliver iron to Pseudomonas, PA14 was co-cultured with *K. aeruginosa* in M9 minimal media with or without supplementation with iron or Pf4 phage. Total Pseudomonas growth and Pseudomonas relative to total bacterial growth (including *K. aeruginosa)* was measured by plating bacteria on selective media (**Fig 4b-c**). We indeed observed that iron associated with Pf phage selectively benefits *P. aeruginosa* in these co-culture systems, as demonstrated by enhanced Pseudomonas growth compared to conditions with M9 media, iron or Pf phage alone. These findings reveal a previously unrecognized mechanism by which a pathogen partners with a virus to selectively enhance iron acquisition and gain a competitive advantage over other microbes.

**Figure 4.**
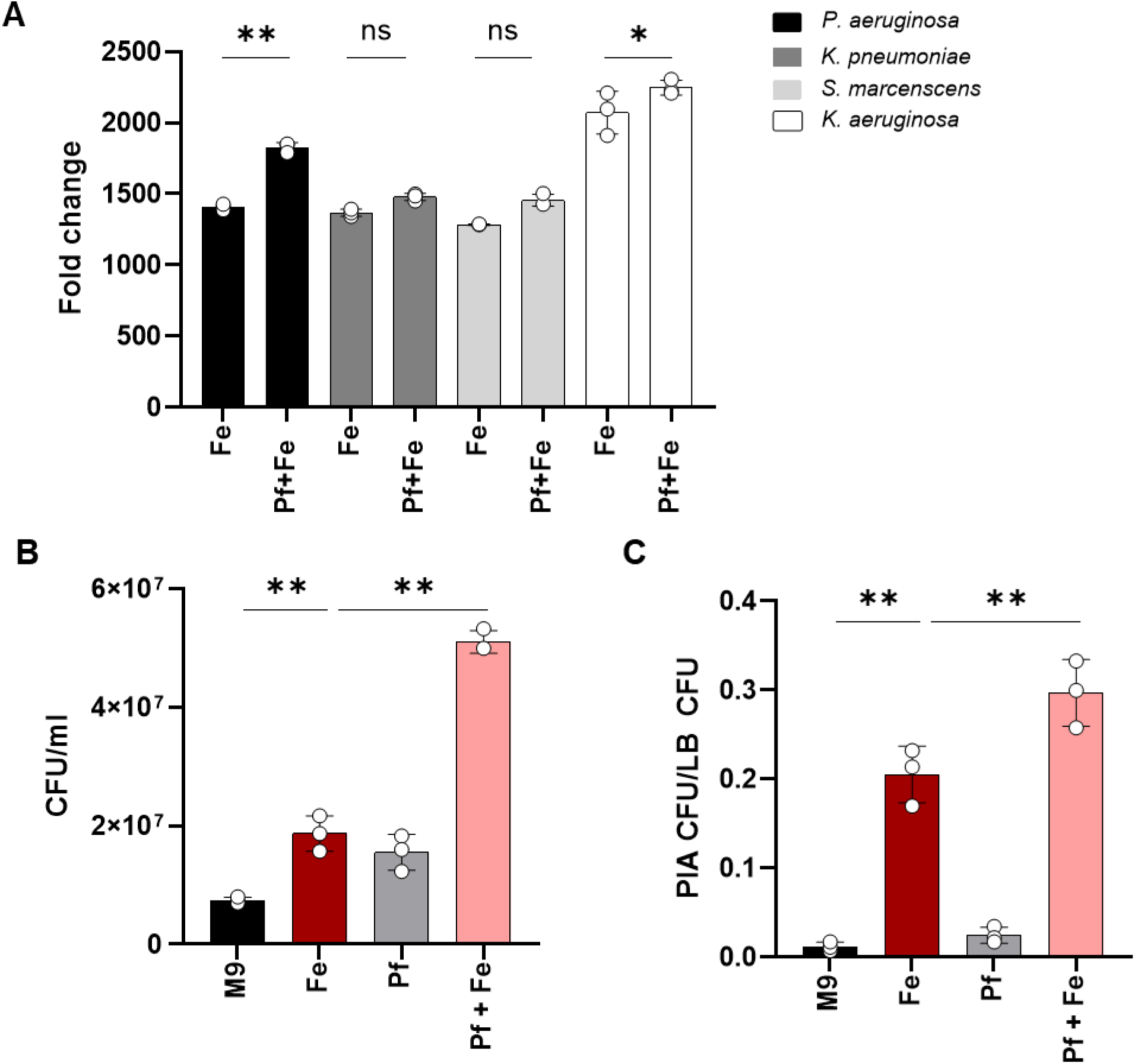
Pf phage selectively promotes iron uptake in *P. aeruginosa*. *P. aeruginosa*, *K. pneumoniae*, *S. marcescens subsp marcescens*, and *K. aeruginosa* are grown in M9 minimal media supplemented with without iron or Pf4 phage. Growth is measured over time and normalized to M9 (A) *P. aeruginosa* and *K. aeruginosa* are grown in M9 minimal media supplemented with without iron or Pf4 phage in 1:1 mixture. Cultures are plated on Pseudomonas Isolation Agar (PIA) and LB agar. B) Total *P. aeruginosa* growth (B) and *P. aeruginosa* normalize to total bacterial growth (C) is quantitated. ** p < 0.01 by one-way ANOVA or Student’s T test.

## Discussion

Iron homeostasis is a critical determinant of *P. aeruginosa* pathogenesis and persistence during chronic infections. Iron functions both as an essential nutrient and as a regulatory signal that coordinates virulence factor expression, biofilm formation, and competitive fitness ^22,37^. Given that chronic *P. aeruginosa* chronic infections remain largely refractory to conventional antimicrobials and contribute to progressive disease with high morbidity, disrupting iron homeostasis has emerged as a promising therapeutic strategy. As such, investigating the specific iron acquisition mechanisms active during chronic infection, and how they shape bacterial persistence and host–pathogen interactions, may reveal critical vulnerabilities that can be exploited to develop more effective, precision-based therapies for chronic *P. aeruginosa* infections.

*P. aeruginosa* has evolved multiple, mechanistically distinct iron acquisition systems, including the endogenous siderophores pyoverdine and pyochelin, uptake of xenosiderophores produced by other microbes, heme utilization pathways, and ferrous iron transport^21^. While this metabolic versatility reflects flexible adaptation to diverse environmental and infection niches, which systems are essential for iron uptake during chronic biofilm infections remains uncertain. The loss of pyoverdine expression clinical isolates from individuals with chronic lung disease suggests alternative iron acquisition strategies, such as heme utilization, is more effective during chronic infection^24,26,39^. Further, exopolysaccharides Psl, Pel and alginate, which form structural components of Pseudomonas biofilms, directly bind iron suggesting a biofilm-specific mechanisms of iron uptake during chronic infections^41^. We now extend these findings by revealing Pf phage, a virus that is highly expressed during chronic infection and forms structural subunits within Pseudomonas biofilms, serves as a unique and previously undescribed mechanism of iron acquisition by bacteria.

Pf phage is an anionic filament which forms crosslinked network polymers in the presence of polycationic metals, including iron. As Pf is a structural subunit within *P. aeruginosa* biofilms, this finding suggests Pf may locally concentrate this essential nutrient to facilitate intracellular uptake. In support of this proposal, we demonstrate Pf phage traps free iron and that Pf-bound iron results in enhanced growth of both *P. aeruginosa* lab stains and clinical isolates, compared to free iron alone. Further, consistent with canonical iron uptake systems, Pf phage is induced during iron-deplete growth conditions driven by PvdS, an iron-starvation ECF sigma factor that induces expression of iron uptake systems.

How Pf phage promotes iron uptake remains unclear. It is possible the Pf binding of ferric iron enhanced its solubility, similar to siderophores, to facilitate uptake. However, Pf also promotes growth in the presence of ferrous iron, which is soluble and does not require chaperon molecules such as siderophores to facilitate uptake. As Pf phage forms crystalline structures that encapsulate *P. aeruginosa*, as alternative possibility is Pf-bound iron is locally more accessible for uptake^43^. Consistent with this possibility, Pf-mediated iron-dependent growth is lost in the absence of type IV pili which is necessary for Pf binding of *P. aeruginosa*.

This study reveals that Pf selectively enhances iron-dependent growth in *P. aeruginosa* through a kin-selective mechanism. Chronic infections are typically polymicrobial, creating intense competition for essential nutrients, particularly iron. To gain a competitive advantage in these environments, bacteria have evolved diverse strategies to appropriate iron from neighboring microbes, such as lysing competitors or hijacking their iron acquisition systems. We previously demonstrated that Pf bacteriophage inhibits the growth of *Aspergillus fumigatus*, an opportunistic fungal pathogen^27^. This inhibitory effect was reversed by iron supplementation, suggesting that Pf limits iron availability to competing fungi^27^. In contrast, in our present study, we find that neither Pf nor Pf-bound iron adversely affects the growth of competing bacterial species, including *Klebsiella pneumoniae, Klebsiella aeruginosa* and *Serratia marcescens*. Rather, Pf-bound iron is selectively utilized by *P. aeruginosa* resulting in enhanced growth and competitive advantage in co-culture conditions. The selective advantage conferred by Pf-mediated iron delivery may contribute to *P. aeruginosa* becoming the dominant bacterial pathogen in polymicrobial chronic infections, particularly in cystic fibrosis airways.

Together, these findings reveal a fundamentally novel mechanism by which a temperate bacteriophage shapes microbial community dynamics and iron homeostasis in chronic Pseudomonas aeruginosa infection. Defining this mechanism has the potential to reveal actionable therapeutic targets, including disruption of Pf production, interference with Pf-mediated iron delivery, or exploitation of Pf-dependent iron uptake for Trojan horse antibiotic strategies, that could inform the development of novel interventions for biofilm-associated infections that are refractory to conventional antimicrobial therapy.

## Methods

### Bacterial strains and culture conditions

See Table 1 for list of strains used in this study. In general, bacteria were prepared as follows. Frozen glycerol stocks were streaked on Luria-Bertani (LB) agar and grown overnight at 37°C. An isolated colony was picked and grown overnight at 37°C in LB medium, under shaking conditions at 37°C.

**Table 1.**

| Strain | Description | Source |
| --- | --- | --- |
| PAO1 | *P. aeruginosa* lab strain, wild-type | (12) |
| PAO1Δpf4 | Deletion of Pf4 prophage from PAO1 | (12) |
| PA14 | *P. aeruginosa* lab strain, wild-type | (<sup>44</sup>) |
| PA14ΔpilC | Deletion of pilC from PA14 resulting in impaired type IV pili function | (<sup>44</sup>) |
| PAO1p | Wild-type parent strain of PAO1 ΔpvdS and PAO1ΔpvdA | Gift from E. Peter Greenberg, University of Washington |
| PAO1ΔpvdS | Deletion of pvdS from PAO1 | Gift from E. Peter Greenberg, University of Washington |
| PAO1ΔpvdA | Deletion of pvdA from PAO1 | Gift from E. Peter Greenberg, University of Washington |
| CPA-53 | *P. aeruginosa* clinical isolate | Gift from Elizabeth Burgener, Children's Hospital Los Angeles |
| CPA-23 | *P. aeruginosa* clinical isolate | Gift from Elizabeth Burgener, Children's Hospital Los Angeles |
| CPA-89 | *P. aeruginosa* clinical isolate | Gift from Elizabeth Burgener, Children's Hospital Los Angeles |
| CPA-16 | *P. aeruginosa* clinical isolate | Gift from Elizabeth Burgener, Children's Hospital Los Angeles |
| *Klebsiella pneumoniae* strain 9633 | Wild-type | ATCC |
| *Serratia marcescens* subsp *marcescens* NCTC 10211 | Wild-type | ATCC |
| *Klebsiella aeruginosa* | Clinical isolate | Gift from Hiutung Chu, University of California, San Diego |

For quantification of Pf phage production under iron-replete or iron-deplete conditions, overnight cultures were diluted to an initial OD₆₀₀ of 0.01 in M9 minimal medium supplemented with ferric chloride at final concentrations ranging from 0.1 to 100 µM, or left un-supplemented as indicated. Following overnight growth, bacterial growth was determined by enumeration of colony-forming units (CFU). Cultures were then centrifuged at 10,000 × g for 5 min to pellet bacteria, and clarified supernatants were passed through 0.2-µm polyethersulfone (PES) filters (MDI Cat. No 1182G76) to remove residual cells. Filtered supernatants were serially diluted and Pf phage titers were quantified by spot plaque assay as described below. Pf production relative to bacterial growth was determined by normalizing PFU to CFU. As a negative control, the isogenic PAO1ΔPf4 strain, in which the Pf4 prophage has been precisely deleted, was included and consistently showed no plaque formation. As an alternative approach, Pf production was assessed by quantification of the Pf4 replicative form (RF) by quantitative PCR (qPCR) as described below.

### Phage purification

This was performed as previously reported (Sweere et al., 2019b). Briefly, *P. aeruginosa* strain PAO1 was grown in LB broth to mid-logarithmic phase and infected with Pf4 phage at a final concentration of 1 × 10¹⁰ PFU/mL. The infected culture was then expanded into 300 mL of LB medium and incubated at 37 °C with shaking for 24 hours. Bacteria were removed by centrifugation at 8,000 × g for 5 minutes, and supernatant was treated with 250 U/µl of Benzonase (Novagen, Cat No. 70746) for 4 hours at 37°C before sterilization. Pf phage were precipitated from the supernatant by adding 0.5 M NaCl and 4% polyethylene glycol (PEG) 8000 (Milipore Sigma, Cat. No. P2139). Phage solutions were incubated overnight at 4°C. Phage were pelleted by centrifugation at 15,000 × g for 30 minutes, and the pellet suspended in sterile TE buffer (pH 8.0). The suspension was centrifuged for 15,000 × g for 20 minutes, and the supernatant was subjected to another round of PEG precipitation. The purified phage pellets were suspended in sterile PBS and dialyzed in 10-kDa molecular weight Slide-A-Lyzer Dialysis Cassette (FisherScientific, Cat. No. 66810) against PBS. Pf concentration was quantified using spot plaque assay, as described below. Working concentration of Pf for growth assays were 1x10^11^ PFU/ml.

### Quantification of Pf phage

Bacteriophage titers were determined by spot plaque assay using the double-layer agar method. Briefly, PAO1ΔPf4 was used as recipient bacteria and was grown overnight in lysogeny broth (LB) at 37°C with shaking, then diluted 1:100 into fresh LB and cultured to mid-logarithmic phase (OD₆₀₀ ≈ 0.4–0.6). Molten top agar consisting of 1% Tryptone (FisherScientific Cat. No. BP1421-2), 1% NaCl (FisherScientific Cat. No. BP 358-212) and 0.5% agar (FisherScientific Cat. No.BP 1423-2) was prepared in advance and maintained at 45–50 °C. PAO1ΔPf4 was added to top agar at a ratio of 1:50 and poured on top of LB agar plates. Once the top-agar overlay had solidified, Pf phage was serially diluted and 20 µL of each dilution was spotted directly onto the surface of the bacterial lawn. Plates were allowed to air-dry until spots were fully absorbed and then incubated inverted at 37°C for 16–24 hours. Pf phage plaques appear as opaque zones of growth inhibition rather than clear plaques. Spots containing countable plaques were used to calculate phage titers, expressed as plaque-forming units per milliliter (PFU/mL), accounting for the dilution factor and volume spotted. Each sample was assayed in technical duplicate or triplicate, and mean titers were reported. Quantification by qPCR was performed as previously described with slight modifications^38^. Bacterial cultures were pelleted at 16,000 × g for 5 minutes, washed 3× in 1× PBS, and treated with DNase at a final concentration of 0.1 mg/mL DNase was heat inactivated and bacterial pellets were lysed by incubating samples at 95 °C for 10 minutes and 1 µl of the lysed bacterial suspension was used as template. qPCR was performed using SsoAdvanced Universal SYBR Green Supermix (BioRad #1725270) on the Roche Lightcycler 480. Pf phage replicative form was determined using the following primers RF For CTT GGC AGG GTG ATT TGG A and RF Rev AGG AAC GCT TCA AAA CCC TA. Normalization to chromosomal copy number was performed using 50S ribosomal protein gene rpIU (rpiU For CAAGGTCCGCATCATCAAGTT and rpiU Rev GGCCCTGACGCTTCATGT). Data was analyzed using the comparative Ct (ΔΔCt) method^40^.

### Growth curve

Bacterial growth curves were performed in 96-well flat-bottom plates (FisherScientific Cat. No FB012931) using M9 minimal medium. Overnight cultures grown in LB broth were diluted 1:1,000 into fresh M9 medium, and 200 µL of the diluted culture was dispensed into each well. Where indicated, vehicle control, ferric chloride at concentrations specified in the figure legends, and/or Pf phage (final concentration 1 × 10¹¹ PFU/mL) were added to designated wells.

Plates were sealed with a Breath-Easy gas permeable sealing membrane (USA Sci Cat. No. 3123-6100) and incubated at 37°C in a Synergy2 Microplate Reader. Bacterial growth was monitored by measuring optical density at 600 nm (OD₆₀₀) at 30-min intervals. Each condition was performed in technical triplicate. Growth data were plotted over time, and the area under the curve (AUC) was calculated to quantify differences in overall growth between conditions.

Bacterial growth and viability were additionally assessed using the BacTiter-Glo microbial cell viability assay (Promega Cat. No. PRG8231), which quantifies intracellular ATP as a proxy for metabolically active bacteria. BacTiter-Glo was used in place of direct colony-forming unit (CFU) enumeration, as it provided a more sensitive and reproducible readout under these conditions. Following completion of the growth curve assay, 100 µL from each well was transferred to a fresh plate and mixed with 100 µL of BacTiter-Glo reagent. Luminescence was measured using a Tecan Spark Multimode Microplate Reader according to the manufacturer’s instructions.

### Imaging PF phage

Purified Pf4 phage, at a concentration of 1 × 10¹⁰ PFU/mL, is incubated with vehicle control or 100 µM iron chloride at room temperature for 30 minutes. After incubation, the mixtures will be diluted 1:10 with deionized water and applied to TEM grids following standard procedures. Briefly, carbon-coated copper grids (300–400 mesh; Cat. No. F300-Cu) is plasma treated, after which 5 µL of each sample is be loaded onto individual grids and negatively stained with 1% uranyl acetate prepared in deionized water. The grids are air-dry at room temperature and subsequently imaged using a JEOL JEM-1400 transmission electron microscope.

### Pyoverdine quantification

Pyoverdine was measured as previously described with slight modifications^42^. The fluorescence of a 100 µl of filtered supernatant was measured (excitation 380, emission 447) and normalized to bacterial CFU.

### Competitive co-culture

PA14 or K. aeruginosa was grown overnight in LB broth and subculture 1:1 in M9 media with final OD₆₀₀ 0.01. 200 µl was aliquoted per well in 96-well flat-bottom plates (FisherScientific Cat. No FB012931) and ferric chloride (final concentration 100 µM) and/or Pf4 phage (final concentration 1x10^11^PFU/ml) were added to designated wells. Plates were incubated at 37°C and aliquots were collected at various timepoints for CFU quantification on both LB agar and Pseudomonas Isolation Agar (PIA; Bio-world Cat. No. 30620067) which were then incubated overnight at 37 °C and CFU formation quantitated.

### Quantification and statistical analysis

All statistical analyses were performed using GraphPad Prism (GraphPad Software, Inc., La Jolla, CA). For bacterial growth curve experiments, the area under the curve (AUC) of OD₆₀₀ measurements over time was calculated and used for statistical comparisons between conditions. Two-tailed unpaired Student’s *t* tests were used for comparisons between two groups, and one-way analysis of variance (ANOVA) with appropriate post hoc tests was used for comparisons among multiple groups. Data are presented as means ± standard error of the mean (SEM) or standard deviation (SD), as indicated in the figure legends. Statistical significance was defined as *p* < 0.05. Additional statistical details, including sample size and specific tests used, are provided in the figure legends.

## Acknowledgements

A.K. discloses support from the Cystic Fibrosis Foundation Fourth Year Clinical Fellowship, Doris Duke Charitable Foundation Physician-Scientist Fellowship and National Institutes of Health grant K08AI190313-01. P.L.B. discloses support from National Institutes of Health grants R01HL148184-01, R01AI12492093, K24 AI166718, and 1R01AI182349-01A1, the Stanford Woods Institute for the Environment, the Stanford-Coulter Translational Research Program, Bio-X, Stanford SPARK, and the Stanford Innovative Medicines Accelerator.

## References

1. Blanchette, C.M., Noone, J.M., Stone, G., Zacherle, E., Patel, R.P., Howden, R., and Mapel, D. (2017). Healthcare Cost and Utilization before and after Diagnosis of Pseudomonas aeruginosa among Patients with Non-Cystic Fibrosis Bronchiectasis in the U.S. Med. Sci. Basel Switz. 5, E20. 10.3390/medsci5040020.

2. Emerson, J., Rosenfeld, M., McNamara, S., Ramsey, B., and Gibson, R.L. (2002). Pseudomonas aeruginosa and other predictors of mortality and morbidity in young children with cystic fibrosis. Pediatr. Pulmonol. 34, 91–100. 10.1002/ppul.10127.

3. Bhagirath, A.Y., Li, Y., Somayajula, D., Dadashi, M., Badr, S., and Duan, K. (2016). Cystic fibrosis lung environment and Pseudomonas aeruginosa infection. BMC Pulm. Med. 16, 174. 10.1186/s12890-016-0339-5.

4. Kosorok, M.R., Zeng, L., West, S.E., Rock, M.J., Splaingard, M.L., Laxova, A., Green, C.G., Collins, J., and Farrell, P.M. (2001). Acceleration of lung disease in children with cystic fibrosis after Pseudomonas aeruginosa acquisition. Pediatr. Pulmonol. 32, 277–287. 10.1002/ppul.2009.abs.

5. Maurice, N.M., Bedi, B., and Sadikot, R.T. (2018). Pseudomonas aeruginosa Biofilms: Host Response and Clinical Implications in Lung Infections. Am. J. Respir. Cell Mol. Biol. 58, 428–439. 10.1165/rcmb.2017-0321TR.

6. Sharma, D., Misba, L., and Khan, A.U. (2019). Antibiotics versus biofilm: an emerging battleground in microbial communities. Antimicrob. Resist. Infect. Control 8, 76. 10.1186/s13756-019-0533-3.

7. Blaznik, M., Volk, M., Kraigher, B., Calonge-Sanz, A., Barco-García, G., Stopar, D., and Dogsa, I. (2025). Biofilm structure as a key factor in antibiotic tolerance: insights from Bacillus subtilis model systems. Npj Biofilms Microbiomes 11, 232. 10.1038/s41522-025-00864-x.

8. Colque, C.A., Albarracín Orio, A.G., Feliziani, S., Marvig, R.L., Tobares, A.R., Johansen, H.K., Molin, S., and Smania, A.M. (2020). Hypermutator Pseudomonas aeruginosa Exploits Multiple Genetic Pathways To Develop Multidrug Resistance during Long-Term Infections in the Airways of Cystic Fibrosis Patients. Antimicrob. Agents Chemother. 64, 10.1128/aac.02142-19. 10.1128/aac.02142-19.

9. Ciofu, O., Mandsberg, L.F., Wang, H., and Høiby, N. (2012). Phenotypes selected during chronic lung infection in cystic fibrosis patients: implications for the treatment of Pseudomonas aeruginosa biofilm infections. FEMS Immunol. Med. Microbiol. 65, 215–225. 10.1111/j.1574-695X.2012.00983.x.

10. Secor, P.R., Burgener, E.B., Kinnersley, M., Jennings, L.K., Roman-Cruz, V., Popescu, M., Van Belleghem, J.D., Haddock, N., Copeland, C., Michaels, L.A., et al. (2020). Pf Bacteriophage and Their Impact on Pseudomonas Virulence, Mammalian Immunity, and Chronic Infections. Front. Immunol. 11, 244. 10.3389/fimmu.2020.00244.

11. Sweere, J.M., Van Belleghem, J.D., Ishak, H., Bach, M.S., Popescu, M., Sunkari, V., Kaber, G., Manasherob, R., Suh, G.A., Cao, X., et al. (2019). Bacteriophage trigger antiviral immunity and prevent clearance of bacterial infection. Science 363, eaat9691. 10.1126/science.aat9691.

12. Bach, M.S., de Vries, C.R., Khosravi, A., Sweere, J.M., Popescu, M.C., Chen, Q., Demirdjian, S., Hargil, A., Van Belleghem, J.D., Kaber, G., et al. (2022). Filamentous bacteriophage delays healing of Pseudomonas-infected wounds. Cell Rep. Med. 3, 100656. 10.1016/j.xcrm.2022.100656.

13. Rice, S.A., Tan, C.H., Mikkelsen, P.J., Kung, V., Woo, J., Tay, M., Hauser, A., McDougald, D., Webb, J.S., and Kjelleberg, S. (2009). The biofilm life cycle and virulence of Pseudomonas aeruginosa are dependent on a filamentous prophage. ISME J. 3, 271–282. 10.1038/ismej.2008.109.

14. Burgener, E.B., Sweere, J.M., Bach, M.S., Secor, P.R., Haddock, N., Jennings, L.K., Marvig, R.L., Johansen, H.K., Rossi, E., Cao, X., et al. (2019). Filamentous bacteriophages are associated with chronic Pseudomonas lung infections and antibiotic resistance in cystic fibrosis. Sci. Transl. Med. 11, eaau9748. 10.1126/scitranslmed.aau9748.

15. Burgener, E.B., Gupta, A., Nakano, K., Gibbs, S.L., Sommers, M.E., Khosravi, A., Bach, M.S., Dunn, C., Spano, J., Secor, P.R., et al. (2024). Pf bacteriophage is associated with decline in lung function in a longitudinal cohort of patients with cystic fibrosis and Pseudomonas airway infection. J. Cyst. Fibros. Off. J. Eur. Cyst. Fibros. Soc., S1569-1993(24)01786-7. 10.1016/j.jcf.2024.09.018.

16. Chen, Q., Cai, P., Chang, T.H.W., Burgener, E., Kratochvil, M.J., Gupta, A., Hargill, A., Secor, P.R., Nielsen, J.E., Barron, A.E., et al. (2024). Pf bacteriophages hinder sputum antibiotic diffusion via electrostatic binding. Sci. Adv. 10, eadl5576. 10.1126/sciadv.adl5576.

17. Secor, P.R., Sweere, J.M., Michaels, L.A., Malkovskiy, A.V., Lazzareschi, D., Katznelson, E., Rajadas, J., Birnbaum, M.E., Arrigoni, A., Braun, K.R., et al. (2015). Filamentous Bacteriophage Promote Biofilm Assembly and Function. Cell Host Microbe 18, 549–559. 10.1016/j.chom.2015.10.013.

18. Secor, P.R., Michaels, L.A., Ratjen, A., Jennings, L.K., and Singh, P.K. (2018). Entropically driven aggregation of bacteria by host polymers promotes antibiotic tolerance in Pseudomonas aeruginosa. Proc. Natl. Acad. Sci. 115, 10780–10785. 10.1073/pnas.1806005115.

19. Tang, J.X., Janmey, P.A., Lyubartsev, A., and Nordenskiöld, L. (2002). Metal Ion-Induced Lateral Aggregation of Filamentous Viruses fd and M13. Biophys. J. 83, 566–581. 10.1016/S0006-3495(02)75192-8.

20. Janmey, P.A., Slochower, D.R., Wang, Y.-H., Wen, Q., and Cēbers, A. (2014). Polyelectrolyte properties of filamentous biopolymers and their consequences in biological fluids. Soft Matter 10, 1439–1449. 10.1039/C3SM50854D.

21. Minandri, F., Imperi, F., Frangipani, E., Bonchi, C., Visaggio, D., Facchini, M., Pasquali, P., Bragonzi, A., and Visca, P. (2016). Role of Iron Uptake Systems in Pseudomonas aeruginosa Virulence and Airway Infection. Infect. Immun. 84, 2324–2335. 10.1128/IAI.00098-16.

22. Banin, E., Vasil, M.L., and Greenberg, E.P. (2005). Iron and Pseudomonas aeruginosa biofilm formation. Proc. Natl. Acad. Sci. 102, 11076–11081. 10.1073/pnas.0504266102.

23. Cornelis, P., Tahrioui, A., Lesouhaitier, O., Bouffartigues, E., Feuilloley, M., Baysse, C., and Chevalier, S. (2022). High affinity iron uptake by pyoverdine in Pseudomonas aeruginosa involves multiple regulators besides Fur, PvdS, and FpvI. BioMetals. 10.1007/s10534-022-00369-6.

24. Cornelis, P., and Dingemans, J. (2013). Pseudomonas aeruginosa adapts its iron uptake strategies in function of the type of infections. Front. Cell. Infect. Microbiol. 3, 75. 10.3389/fcimb.2013.00075.

25. Konings, A.F., Martin, L.W., Sharples, K.J., Roddam, L.F., Latham, R., Reid, D.W., and Lamont, I.L. (2013). Pseudomonas aeruginosa Uses Multiple Pathways To Acquire Iron during Chronic Infection in Cystic Fibrosis Lungs. Infect. Immun. 81, 2697–2704. 10.1128/IAI.00418-13.

26. Nguyen, A.T., O’Neill, M.J., Watts, A.M., Robson, C.L., Lamont, I.L., Wilks, A., and Oglesby-Sherrouse, A.G. (2014). Adaptation of Iron Homeostasis Pathways by a Pseudomonas aeruginosa Pyoverdine Mutant in the Cystic Fibrosis Lung. J. Bacteriol. 10.1128/JB.01491-14.

27. Penner, J.C., Ferreira, J.A.G., Secor, P.R., Sweere, J.M., Birukova, M.K., Joubert, L.-M., Haagensen, J.A.J., Garcia, O., Malkovskiy, A.V., Kaber, G., et al. (2016). Pf4 bacteriophage produced by Pseudomonas aeruginosa inhibits Aspergillus fumigatus metabolism via iron sequestration. Microbiol. Read. Engl. 162, 1583–1594. 10.1099/mic.0.000344.

28. Hunter, R.C., Asfour, F., Dingemans, J., Osuna, B.L., Samad, T., Malfroot, A., Cornelis, P., and Newman, D.K. (2013). Ferrous iron is a significant component of bioavailable iron in cystic fibrosis airways. mBio 4, e00557–13. 10.1128/mBio.00557-13.

29. Whiteley, M., Bangera, M.G., Bumgarner, R.E., Parsek, M.R., Teitzel, G.M., Lory, S., and Greenberg, E.P. (2001). Gene expression in Pseudomonas aeruginosa biofilms. Nature 413, 860–864. 10.1038/35101627.

30. Guo, Y., Tang, K., Sit, B., Gu, J., Chen, R., Shao, X., Lin, S., Huang, Z., Nie, Z., Lin, J., et al. (2024). Control of lysogeny and antiphage defense by a prophage-encoded kinase-phosphatase module. Nat. Commun. 15, 7244. 10.1038/s41467-024-51617-x.

31. Ismail, M.H., Michie, K.A., Goh, Y.F., Noorian, P., Kjelleberg, S., Duggin, I.G., McDougald, D., and Rice, S.A. (2021). The Repressor C Protein, Pf4r, Controls Superinfection of Pseudomonas aeruginosa PAO1 by the Pf4 Filamentous Phage and Regulates Host Gene Expression. Viruses 13, 1614. 10.3390/v13081614.

32. Ochsner, U.A., Wilderman, P.J., Vasil, A.I., and Vasil, M.L. (2002). GeneChip® expression analysis of the iron starvation response in Pseudomonas aeruginosa: identification of novel pyoverdine biosynthesis genes. Mol. Microbiol. 45, 1277–1287. 10.1046/j.1365-2958.2002.03084.x.

33. Rédly, G.A., and Poole, K. (2005). FpvIR Control of fpvA Ferric Pyoverdine Receptor Gene Expression in Pseudomonas aeruginosa: Demonstration of an Interaction between FpvI and FpvR and Identification of Mutations in Each Compromising This Interaction. J. Bacteriol. 187, 5648–5657. 10.1128/JB.187.16.5648-5657.2005.

34. Rédly, G.A., and Poole, K. (2005). FpvIR Control of fpvA Ferric Pyoverdine Receptor Gene Expression in Pseudomonas aeruginosa: Demonstration of an Interaction between FpvI and FpvR and Identification of Mutations in Each Compromising This Interaction. J. Bacteriol. 187, 5648–5657. 10.1128/jb.187.16.5648-5657.2005.

35. Salsgiver, E.L., Fink, A.K., Knapp, E.A., LiPuma, J.J., Olivier, K.N., Marshall, B.C., and Saiman, L. (2016). Changing Epidemiology of the Respiratory Bacteriology of Patients With Cystic Fibrosis. Chest 149, 390–400. 10.1378/chest.15-0676.

36. Parkins, M.D., Somayaji, R., and Waters, V.J. (2018). Epidemiology, Biology, and Impact of Clonal Pseudomonas aeruginosa Infections in Cystic Fibrosis. Clin. Microbiol. Rev. 31, e00019–18. 10.1128/CMR.00019-18.

37. Smith, D.J., Lamont, I.L., Anderson, G.J., and Reid, D.W. (2013). Targeting iron uptake to control Pseudomonas aeruginosa infections in cystic fibrosis. Eur. Respir. J. 42, 1723–1736. 10.1183/09031936.00124012.

38. Schmidt, A.K., Schwartzkopf, C.M., Pourtois, J.D., Burgener, E.B., Faith, D.R., Joyce, A., Lamma, T., Kumar, G., Bollyky, P.L., and Secor, P.R. (2024). Targeted deletion of Pf prophages from diverse Pseudomonas aeruginosa isolates has differential impacts on quorum sensing and virulence traits. J. Bacteriol. 206, e00402–23. 10.1128/jb.00402-23.

39. Marvig, R.L., Damkiær, S., Khademi, S.M.H., Markussen, T.M., Molin, S., and Jelsbak, L. (2014). Within-host evolution of Pseudomonas aeruginosa reveals adaptation toward iron acquisition from hemoglobin. mBio 5, e00966–00914. 10.1128/mBio.00966-14.

40. Schmittgen, T.D., and Livak, K.J. (2008). Analyzing real-time PCR data by the comparative C(T) method. Nat. Protoc. 3, 1101–1108. 10.1038/nprot.2008.73.

41. Yu, S., Wei, Q., Zhao, T., Guo, Y., and Ma, L.Z. (2016). A Survival Strategy for Pseudomonas aeruginosa That Uses Exopolysaccharides To Sequester and Store Iron To Stimulate Psl-Dependent Biofilm Formation. Appl. Environ. Microbiol. 82, 6403–6413. 10.1128/AEM.01307-16.

42. Voulhoux, R., Filloux, A., and Schalk, I.J. (2006). Pyoverdine-Mediated Iron Uptake in Pseudomonas aeruginosa: the Tat System Is Required for PvdN but Not for FpvA Transport. J. Bacteriol. 188, 3317–3323. 10.1128/JB.188.9.3317-3323.2006.

43. Tarafder, A.K., von Kügelgen, A., Mellul, A.J., Schulze, U., Aarts, D.G.A.L., and Bharat, T.A.M. (2020). Phage liquid crystalline droplets form occlusive sheaths that encapsulate and protect infectious rod-shaped bacteria. Proc. Natl. Acad. Sci. U. S. A. 117, 4724–4731. 10.1073/pnas.1917726117.

44. Kim, M.K., Chen, Q., Echterhof, A., McBride, R.C., Pennetzdorfer, N., Banaei, N., Burgener, E.B., Milla, C.E., and Bollyky, P.L. (2024). A Blueprint for Broadly Effective Bacteriophage Therapy Against Bacterial Infections. Preprint at bioRxiv, 10.1101/2024.04.20.590411 10.1101/2024.04.20.590411.

