## Supplemental Figures for "Phage-Mediated Iron Acquisition by *Pseudomonas aeruginosa*"

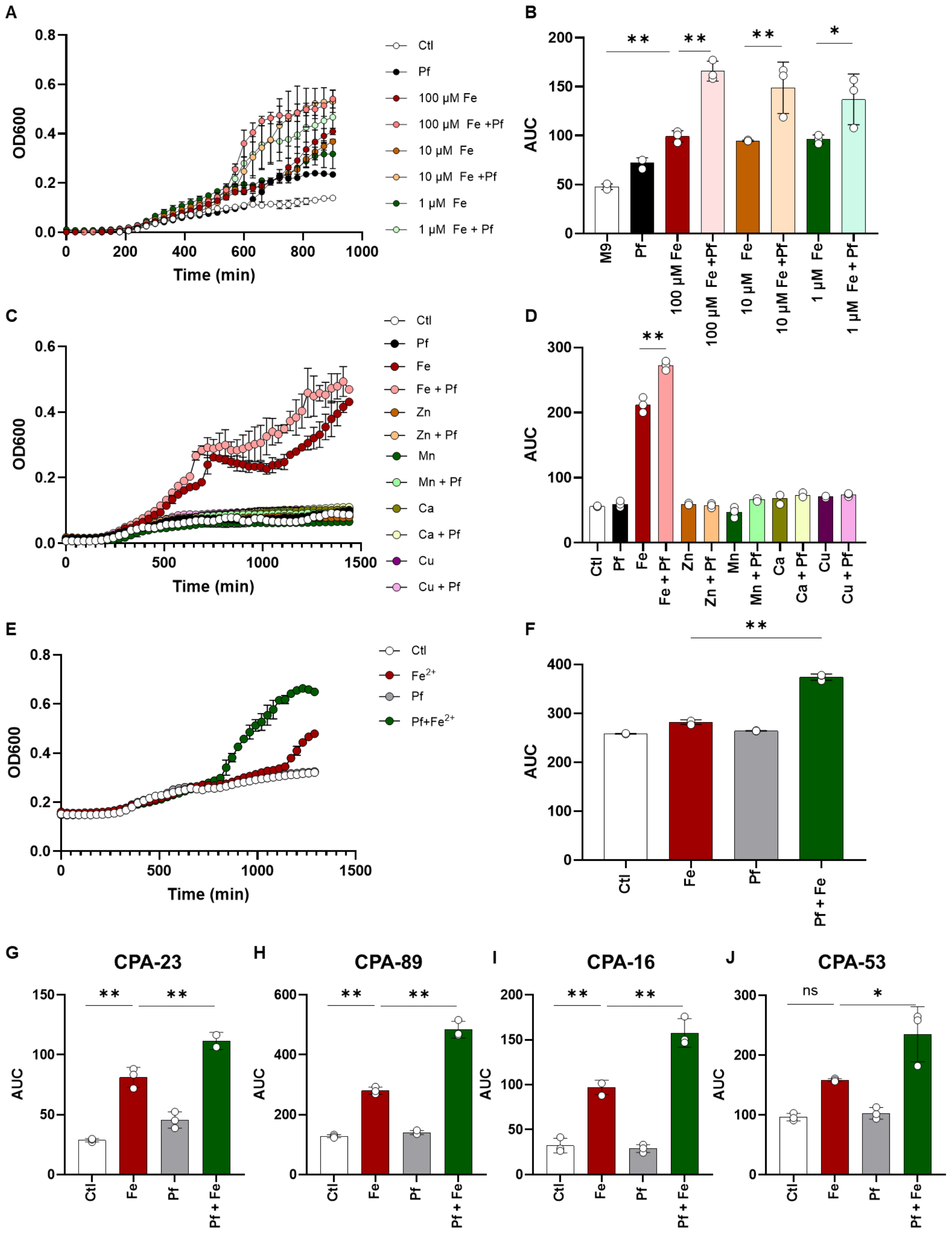

A
B
C
D
E
F
G
H
I
J
OD600
Time (min)
AUC
Ctl
Pf
100 μM Fe
100 μM Fe +Pf
10 μM Fe
10 μM Fe +Pf
1 μM Fe
1 μM Fe + Pf
M9
Fe
Fe + Pf
Zn
Zn + Pf
Mn
Mn + Pf
Ca
Ca + Pf
Cu
Cu + Pf
Fe2+
Pf+Fe2+
Pf + Fe
CPA-23
CPA-89
CPA-16
CPA-53
ns

**Supplementary Figure 2. Pf selectively promotes iron-dependent growth.** PA14 is grown in M9 minimal media (Control, Ctl) or with 1-100 µM ferric chloride with or without Pf4 ($1x10^{11}$ PFU/ml final concentration). Growth is measured over time (A) and area under the curve (AUC) is quantitated (B). PA14 is grown in M9 minimal media (Ctl) or in M9 supplemented with iron (Fe), zinc (Zn), manganese (Mn), calcium (Ca), copper (Cu) or Pf4 phage (Pf). Growth is measured over time (C) and area under the curve (AUC) is quantitated (D). PA14 is grown in M9 minimal media supplemented with 2 mM ascorbic acid alone (Ctl) or supplemented with Pf phage or 100 µM ferric chloride, which in the presence of ascorbic acid is reduced to ferrous iron. Growth and AUC is measured over time (E-F). Clinical *P. aeruginosa* strains CPA-23 (G), CPA-89 (H), CPA-16 (I), CPA-53 (J) are grown in M9 media (Ctl) with or without iron or Pf supplementation as above and growth is measured over time and quantitated as AUC. * $p < 0.05$ ** $p < 0.01$ by one-way ANOVA.

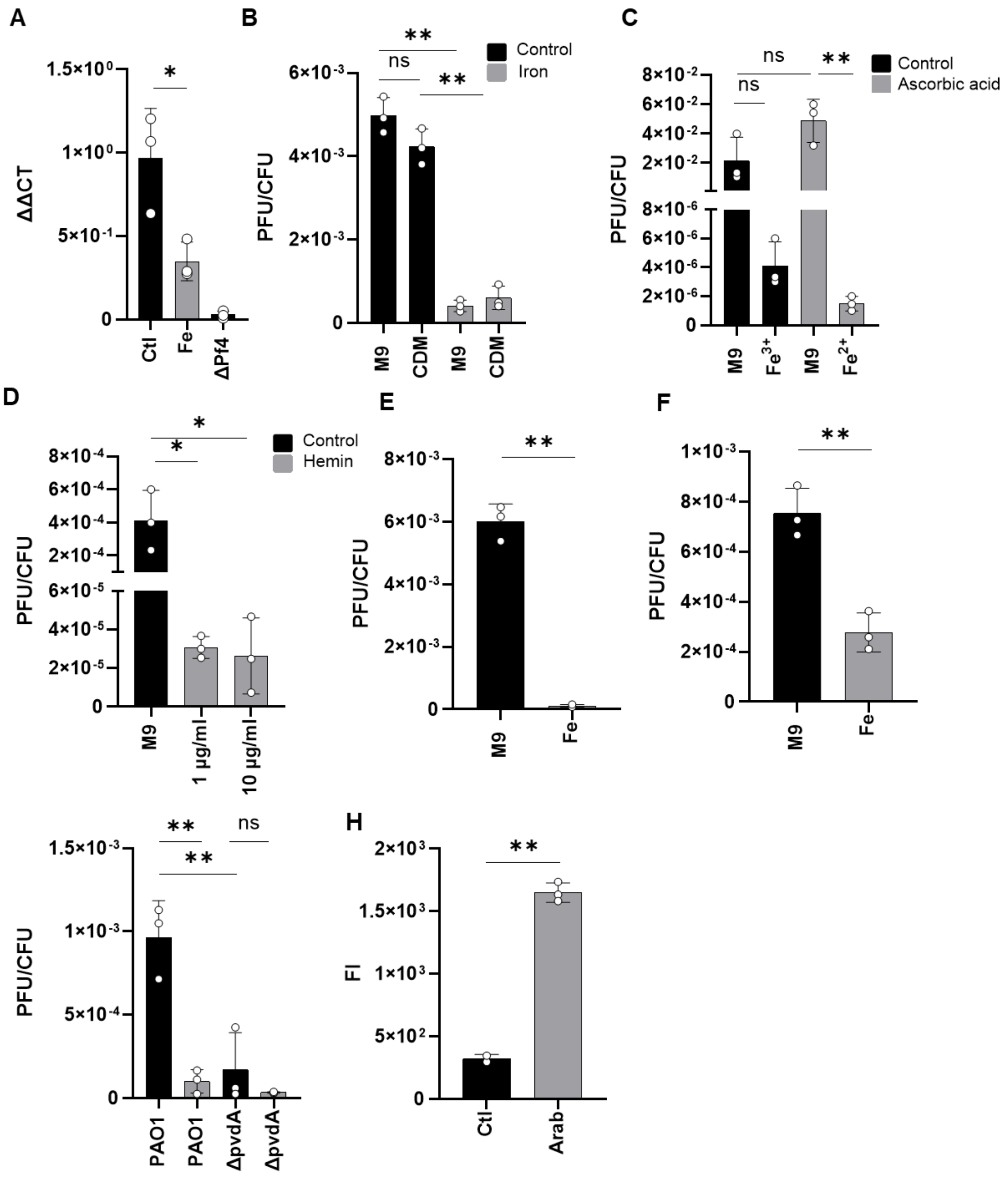


**Supplementary Figure 3. Iron inhibits Pf expression.** A) PAO1 is grown in M9 minimal media without iron supplementation (Ctl) or with 100 μM ferric chloride. Pf replicative form (RF) is quantitated by qPCR. PAO1ΔPf4 is included as a negative control. B) PAO1 is grown in M9 minimal media or chemically defined media (CDM) composed of M9 supplemented with nickel, copper, manganese and zinc. M9 and CDM is supplemented with or without 100 μM iron chloride and Pf expression relative to bacterial growth is measured. C) PAO1 grown in M9 media with or without supplementation with 100 μM ferric chloride and/or 2mM ascorbic acid to reduce ferric iron to ferrous iron. Pf plaque formation is measured at 24 hours and normalized to bacterial growth. D) PAO1 grown in M9 with or without hemin supplementation and Pf expression relative to bacterial growth reported. E) PAO1 biofilms grown in static culture in M9 media (black) or M9 media supplemented with 100 µM ferric chloride (grey). Pf relative to total bacterial CFU is measured. F) CPA-53, a *P. aeruginosa* clinical isolate is grown in M9 media (black) or M9 media supplemented with 100 µM ferric chloride (grey). Pf relative to total bacterial CFU is measured. G) PAO1 or PAO1ΔpvdA grown in M9 minimal media with or without 100 µM ferric chloride supplementation and Pf expression relative to bacterial growth is measured at 24 hours. H) PAO1 containing pvdS cloned under a arabinose (Arab)-inducible promoter treated with vehicle control or 0.5% arabinose and pyoverdine expression is measured. * $p < 0.05$ ** $p < 0.01$ by one-way ANOVA or Student's T-test.

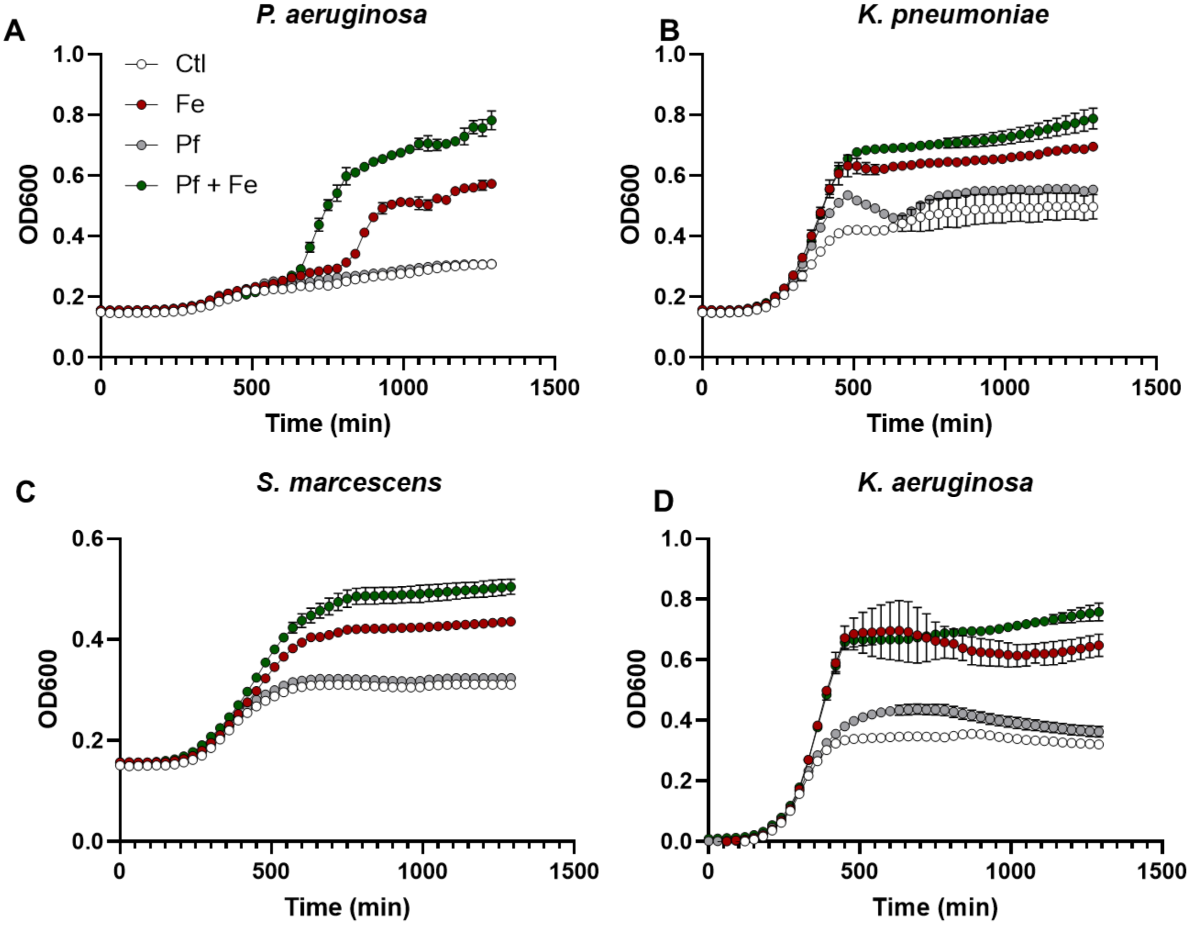


**Supplementary Figure 4. Limited Pf-mediated iron growth in non-Pseudomonas strains.** A) *P. aeruginosa* PA14, B) *K. pneumoniae, C) S. marcescens subspecies marcescens* and D) *K. aeruginosa* are grown in M9 media with or without supplementation with 100 µM ferric chloride or $1x10^{11}$ PFU/ml Pf4 phage and growth is measured over time. Data is representative of three independent experiments.
